# CAR-NK Cells Effectively Target the D614 and G614 SARS-CoV-2-infected Cells

**DOI:** 10.1101/2021.01.14.426742

**Authors:** Minh Ma, Saiaditya Badeti, Chih-Hsiung Chen, Abraham Pinter, Qingkui Jiang, Lanbo Shi, Renping Zhou, Huanbin Xu, Qingsheng Li, William Gause, Dongfang Liu

## Abstract

Severe Acute Respiratory Syndrome Coronavirus 2 (SARS-CoV-2) is highly contagious presenting a significant public health issue. Current therapies used to treat coronavirus disease 2019 (COVID-19) include monoclonal antibody cocktail, convalescent plasma, antivirals, immunomodulators, and anticoagulants, though the current therapeutic options remain limited and expensive. The vaccines from Pfizer and Moderna have recently been authorized for emergency use, which are invaluable for the prevention of SARS-CoV-2 infection. However, their long-term side effects are not yet to be documented, and populations with immunocompromised conditions (e.g., organ-transplantation and immunodeficient patients) may not be able to mount an effective immune response. In addition, there are concerns that wide-scale immunity to SARS-CoV-2 may introduce immune pressure that could select for escape mutants to the existing vaccines and monoclonal antibody therapies. Emerging evidence has shown that chimeric antigen receptor (CAR)- natural killer (NK) immunotherapy has potent antitumor response in hematologic cancers with minimal adverse effects in recent studies, however, the potentials of CAR-NK cells in preventing and treating severe cases of COVID-19 has not yet been fully exploited. Here, we improve upon a novel approach for the generation of CAR-NK cells for targeting SARS-CoV-2 and its D614G mutant. CAR-NK cells were generated using the scFv domain of S309 (henceforward, S309-CAR-NK), a SARS-CoV and SARS-CoV-2 neutralizing antibody that targets the highly conserved region of SARS-CoV-2 spike (S) glycoprotein, therefore would be more likely to recognize different variants of SARS-CoV-2 isolates. S309-CAR-NK cells can specifically bind to pseudotyped SARS-CoV-2 virus and its D614G mutant. Furthermore, S309-CAR-NK cells can specifically kill target cells expressing SARS-CoV-2 S protein *in vitro* and show superior killing activity and cytokine production, compared to that of the recently published CR3022-CAR-NK cells. Thus, these results pave the way for generating ‘off-the-shelf’ S309-CAR-NK cells for treatment in high-risk individuals as well as provide an alternative strategy for patients unresponsive to current vaccines.

## Introduction

SARS-CoV-2 is highly transmissible and has thus far infected millions of people worldwide, approximately one-fifth of the total reported cases are in the US (1). Since February 2020, the frequency of the SARS-CoV-2 D614G variant has increased significantly and has become the dominant variant over the course of the pandemic (2). The D614G variant is shown to associate with higher viral loads in patients, though there is currently insufficient scientific evidence showing the effect of D614G mutation in increased infectivity and transmissibility (3, 4). The disease caused by SARS-CoV-2 (i.e., COVID-19) presents severe symptoms including pneumonia, acute respiratory distress syndrome (5), neurological symptoms, organ failure, and death. More importantly, severe COVID-19 patients may experience dysregulation of an appropriate immune response, characterized by lymphopenia (6), high neutrophil levels in peripheral blood (7), and increased pro-inflammatory cytokines and chemokines (8). While repurposed therapeutics such as monoclonal antibody cocktails, convalescent plasma, and dexamethasone have shown promising results in treating COVID-19 (9). The Pfizer/BioNTech (NCT04368728) and Moderna (NCT04283461) vaccines were recently approved by the FDA for emergency use (10, 11), however, their long-term effects have not yet been documented (12-14). Furthermore, there is accumulating evidence that viral variants that develop relative resistance to the current vaccines and antibody treatments are becoming more common. There could also be subsets of patients who may not be responsive to the vaccines (e.g., people with immune-compromising conditions), and therefore there is a need for an alternative protection or treatment strategy.

Multiple sources of NK cells are found in cord blood (CB), peripheral blood (PB), bone marrow (BM), human embryonic stem cells (ESCs), and induced pluripotent stem cells (iPSC) (15, 16). NK cells isolated from these sources can be further modified to express a chimeric antigen receptor (CAR) for treating a variety of cancer and infectious diseases (17). Recent preclinical studies of CAR-NK in cancer immunotherapy show several advantages of CAR-NK cells over CAR-T cells in clinical safety. For instance, CAR-NK cells do not present additional risk for the development of severe graft-versus-host-disease (GVHD) (18). More importantly, CAR-NK cells are associated with reduced host cytotoxicity compared to CAR-T cells. Specifically, NK cells are less likely to induce cytokine release syndrome (CRS) that could potentially exacerbate COVID-19 symptoms in severe patients (19). Additionally, CAR-NK cells have potential to be developed as a ‘off-the-shelf’ CAR product in the near future (18, 20, 21). Given these aforementioned reasons, NK and CAR-NK cell-based immunotherapeutics have been rapidly developed for COVID-19 treatment. Specifically, adoptive transfer of monocytes or NK cells (NCT04375176, NCT04280224, and NCT04365101) and the universal ‘off-the-shelf’ NKG2D-ACE2 CAR-NK cells (NCT04324996) expanded from CB are currently being studied in clinical trials for COVID-19.

Previous studies show that the genome sequence of SARS-CoV-2 is 79.6% identical to that of SARS-CoV (22). Similar to SARS-CoV, the Spike protein expressed on the surface of SARS-CoV-2 binds to the angiotensin-converting enzyme-2 (ACE2) receptor and facilitates virus entry (22, 23). Several neutralizing antibodies were isolated from memory B cells of convalescent SARS patients which possess cross-reactivity for SARS-CoV-2. One such antibody, named S309, potently neutralizes both pseudotyped SARS-CoV-2 viral particles and wild-type SARS-CoV-2 by binding to both the ‘closed’ and ‘open’ ectodomain trimer conformations of the SARS-CoV-2 Spike glycoprotein (24). In this study, we developed a novel approach for the generation of CAR-NK cells for targeting SARS-CoV-2 using the scFv domain of S309. Here, we show that both the S309-CAR-NK-92MI cell line and primary S309-CAR-NK cells expanded from PB (hereinafter S309-CAR-NK^primary^) cells effectively bind to SARS-CoV-2 pseudovirus and the D614G variant pseudovirus. Moreover, compared to the previously generated Spike-protein-targeting CR3022-CAR-NK cells (25), S309-CAR-NK cells show superior killing activities against target cells (e.g., A549, an epithelial carcinoma derived from a 58 year old Caucasian male with a non-small-cell lung carcinoma (26)) expressing SARS-CoV-2 S protein and mutant S protein (hereinafter, A549-Spike and A549-Spike D614G, respectively) *in vitro*. S309-CAR-NK^primary^ cells show increased productions of TNF-α and IFN-γ when cocultured with A549-Spike and A549-Spike D614G target cells in comparison to CR3022-CAR-NK^primary^. The “4-hour gold standard” chromium release assay further confirmed the superior killing activities of both S309-CAR-NK-92MI cell line and S309-CAR-NK^primary^. These data show that ‘off-the-shelf’ S309-CAR-NK cells may have the potential to prevent SARS-CoV-2 infection, as well as to treat immunocompromised patients or those with comorbidities such as diabetes, cancer, malnutrition, and certain genetic disorders who have been infected with SARS-CoV-2.

## RESULTS

### Generation and characterization of S309-CAR-NK-92MI cells

To develop an NK cell-based immunotherapy for a COVID-19 prevention and treatment, we cloned the scFv domain of S309 into an SFG retroviral vector that contains a human IgG1 hinge and CH2-CH3 domain, CD28 transmembrane (27) domain and intracellular domain, 4-1BB intracellular domain, and CD3zeta intracellular domain (**Fig. 1a**). After construction of S309-CAR, we successfully generated S309-CAR-NK cells in the human NK-92MI cell line (**Fig. 1a**). Briefly, 293T cells were transfected with a combination of plasmids containing S309-CAR in the SFG backbone, retroviral helper plasmids RDF and PegPam3, as previously described (28). The SFG retrovirus particles were then used to transduce NK-92MI cells. After 4-5 days, NK-92MI and S309-CAR cells were stained with CD56 and human IgG (H+L) and the CAR expression was analyzed by flow cytometry. Around 70% of CD56^+^ S309-CAR^+^ NK-92MI cells were observed (**Fig. 1b**). Then, the subsequent S309-CAR positive NK-92 cells were sorted by flow cytometry to achieve high CAR expression levels (**Fig. 1b**). In summary, we have successfully established the surface expression of S309-CAR-NK cells.

**Figure 1:**
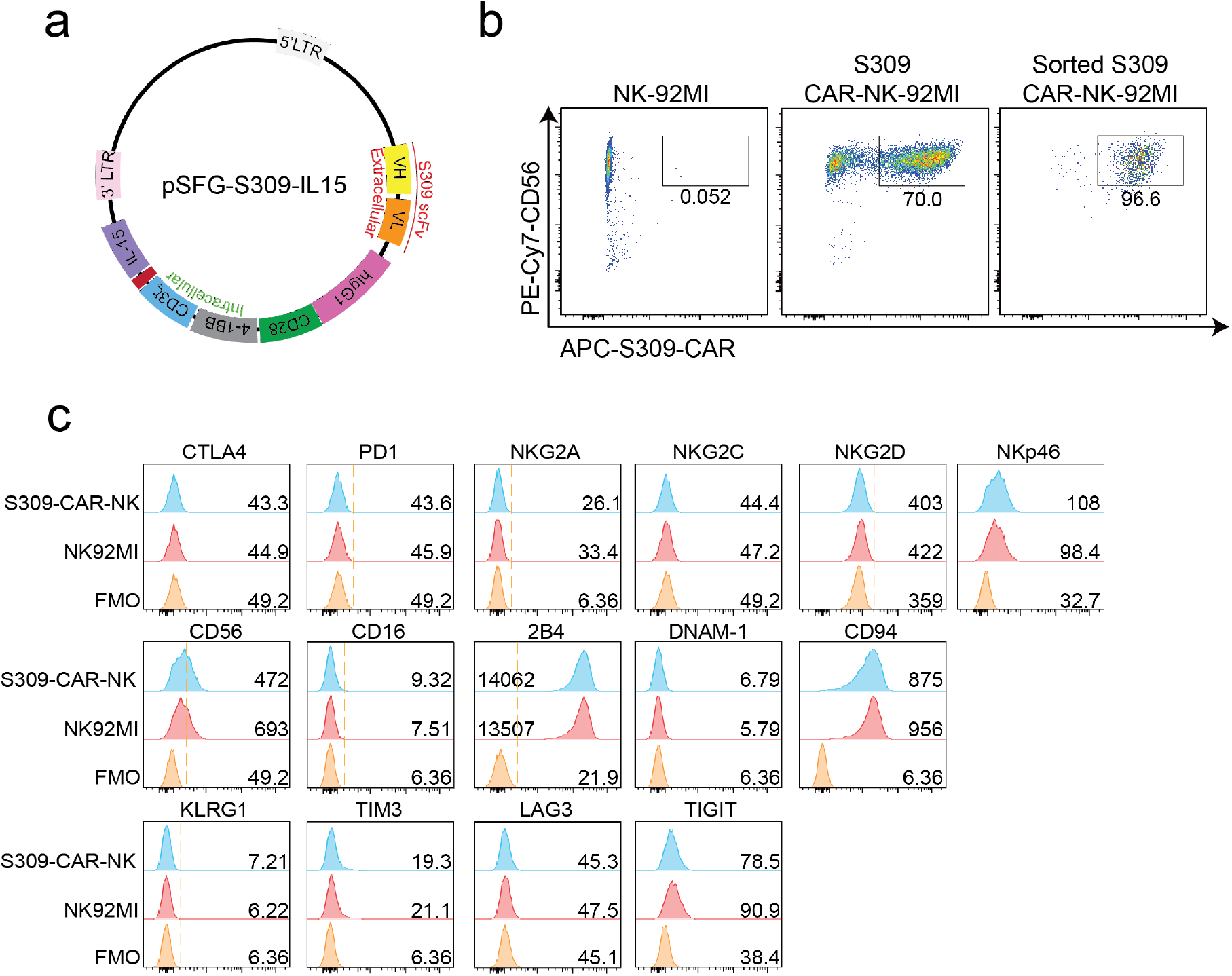
Generation of S309-CAR-NK-92MI cells. **(a**) Plasmid construct of S309-CAR. The SFG retroviral vector contains the S309 single chain antibody fragment (accession code 6WS6 on PDB), a human IgG1 CH2CH3 hinge region, a CD28 transmembrane region, followed by the co-stimulatory CD28, 4-1BB, and the intracellular domain of CD3ζ. (**b**) Determination of S309-CAR-NK expression by flow cytometry. S309-CAR cells were collected and stained with anti-CD56 and CAR F(ab)2 domain [IgG (H+L)] for flow cytometry. The cells were then sorted to achieve a homogenous population of high CAR expression. (**c**) Immunoprofiling of S309-CAR-NK-92MI cells by flow cytometry. S309-CAR and the wildtype NK-92MI cells were stained with antibodies against different immunomodulatory receptors including CTLA4, PD1, NKG2A, NKG2C, NKG2D, NKp46, CD56, CD16, 2B4, DNAM-1, CD94, KLRG1, TIM3, LAG3, and TIGIT.

To characterize S309-CAR-NK-92MI cells, we examined the expressions of several key immunoreceptors on S309-CAR-NK-92MI cells by flow cytometry. These receptors include TIGIT, LAG-3, TIM-3, KLRG1, CTLA-4, PD-1, CD69, CD8A, NKG2C, CD94, DNAM-1, 2B4, NKG2D, NKp46, and CD16 (**Fig. 1c**). Overall, the expressions of these activating and inhibitory receptors are comparable between parental NK-92MI and S309-CAR-NK-92MI cells, indicating the comparable characteristics of NK-92MI at pre- and post-transduction stages.

### S309-CAR-NK cells bind to the immunogenic receptor-binding domain (RBD) domain of SARS-CoV-2 and pseudotyped SARS-CoV-2 S viral particles

After successful establishment of S309-CAR-NK-92MI cells, we then assessed the binding ability of S309-CAR-NK cells to the RBD domain of SARS-CoV-2 S protein. Since S309 neutralizing antibody was isolated from memory B cells of a SARS patient (24), we also included the recombinant His-RBD protein of SARS-CoV as a positive control. S309-CAR-NK-92MI cells and NK-92MI cells were incubated with the His-RBD of SARS-CoV or RBD-SARS-CoV-2 and the resulting complex was then recognized by anti-His and its corresponding fluorophore-conjugated-secondary antibody. Flow cytometry was employed to evaluate the binding efficiency of S309-CAR to the RBD of S protein from either SARS-CoV or SARS-CoV-2. Consistent with results from previous studies (29), S309 recognizes and strongly binds to the RBD of both SARS-CoV and SARS-CoV-2 (**Fig. 2a**). We therefore conclude that S309-CAR-NK-92MI cells can specifically bind to the recombinant His-RBD protein of SARS-CoV and the RBD-SARS-CoV-2.

**Figure 2:**
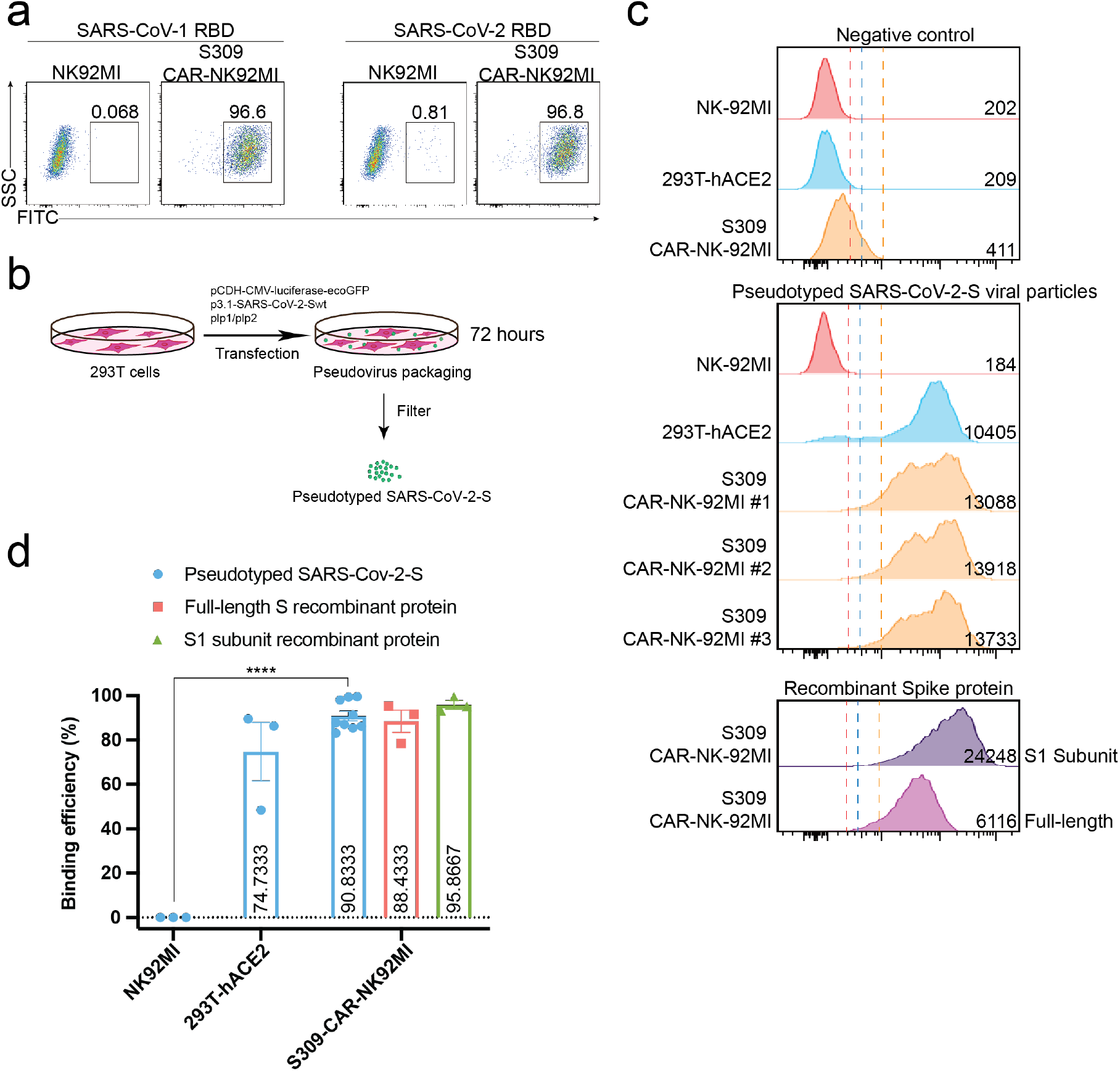
S309-CAR-NK-92MI cells bind to RBD domain of SARS-CoV-2 S protein and pseudotyped SARS-CoV-2 viral particles. (**a**) Representative dot plots showing the efficiency of S309-CAR binding to SARS-CoV-2-RBD. S309-CAR-NK-92MI or parental NK-92MI cells were incubated with the RBD recombinant protein of SARS-CoV-1 or SARS-CoV-2. (**b**) Generation of pseudotyped SARS-CoV-2 viral particles. 293T cells were transfected with various plasmids for 72 hours for the generation of pseudotyped SARS-CoV-2 viral particles. (**c**) Representative histogram showing S309-CAR-NK-92MI binds to the pseudotyped SARS-CoV-2 viral particles. S309-CAR-NK-92MI, parental NK-92MI, or 293T-hACE2 (positive control) cells were incubated with pseudotyped SARS-CoV-2 viral particles, S1 subunit, or full-length S recombinant protein at 37°C for 1 hour. The experimental sample was performed in triplicates with MFI = 13579 ± 251 (a.u.). (**d**) Quantitative data of the binding efficiency of S309-CAR-NK-92MI cells to pseudotyped SARS-CoV-2 viral particles. The experimental sample was performed in triplicates with binding efficiency of over 90%. Data represent mean ± standard error of the mean (SEM) of three independent experiments. Unpaired Student’s *t* test was employed. ****p < 0.0001.

However, the partial RBD domain of SARS-CoV-2 S may not fully reflect the complexity of SARS-CoV-2 viral particles. We therefore evaluated the binding ability of S309-CAR-NK cells to pseudotyped SARS-CoV-2 S viral particles that we generated. Pseudotyped SARS-CoV-2 viral particles were produced by transfecting 293T cells with a combination of pCMV-luciferase-ecoGFP, pcDNA3.1-SARS-CoV-2 Spike, pLP1, and pLP2 plasmids. The supernatant containing pseudotyped SARS-CoV-2 viral particles were used for the pseudovirus binding assay (**Fig. 2b**). To further confirm the presence of pseudotyped SARS-CoV-2 viral particles, we used the collected SARS-CoV-2 pseudovirus to infect 293T-hACE2 cells. After 48-72 hours of incubation, we observed the GFP expression of the infected 293T-hACE2 cells by EVOS florescence microscope (data not shown) and flow cytometry analysis **(Fig. S2)**.

Previous studies show that the RBD of S glycoprotein binds to ACE2 and facilitates SARS-CoV-2 entry (30). Thus, we included 293T-hACE2 in our experiment as a positive control (**Fig. 2c**). Full-length Spike and RBD-containing S1 subunit recombinant proteins were also included as additional control groups (**Fig. 2c & 2d**). To evaluate the binding ability of S309-CAR-NK-92MI to the pseudotyped SARS-CoV-2 S virus, S309-CAR-NK-92MI, NK-92MI or 293T-hACE2 were incubated with SARS-CoV-2 S viral particles, S1 subunit, or full-length S recombinant protein. The complex can be recognized by anti-S1 subunit antibody and its corresponding fluorophore-conjugated secondary antibody. As expected, S309-CAR-NK-92MI cells were able to bind to the pseudotyped SARS-CoV-2 S viral particles with slightly lower binding efficiency than that of recombinant protein groups (**Fig. 2c**). Surprisingly, S309-CAR-NK cells showed a stronger binding efficiency to the pseudotyped SARS-CoV-2 viral particles compared to that of 293T-hACE2 cells, suggesting that S309-CAR-NK-92MI may have superior binding capabilities to the SARS-CoV-2 virus compared to the natural receptor, ACE2 (**Fig. 2d**). In summary, S309-CAR-NK-92MI cells can strongly and specifically bind to the RBD-containing S1 subunit recombinant protein and the full-length pseudotyped SARS-CoV-2 S viral particles.

### S309-CAR-NK cells can be activated by SARS-CoV-2 spike protein RBD expressing target cells and specifically kill their susceptible target cells

After successful generation of S309-CAR-NK cells and demonstration of binding activities of recombinant His-RBD protein of SARS-CoVs (including SARS-CoV-1 and SARS-CoV-2) and pseudotyped full length S viral particle derived from SARS-CoV-2, we further evaluated whether S309-CAR-NK cells can be activated by target cells expressing SARS-CoV-2 S protein. To test this, we generated two different cell lines expressing the RBD and S proteins using 293T-hACE2 and A549, respectively. For the generation of transient 293T-hACE2-RBD cells, we transfected an RBD encoding plasmid into 293T-hACE2 cells (a commonly used cell line for studying the SARS-CoV-2 virus) (**Fig. 3a**). On average, the transfection efficiencies of RBD proteins on 293T-hACE2 cells were greater than 90% as determined by flow cytometry (**Fig. 3b**) and confocal microscopy (data not shown). For the generation of the stable A549-Spike cell line (**Table S1**), we transfected 293T cells with a combination of plasmids containing RDF, Pegpam3, and SARS-CoV-2 S in SFG backbone to produce SARS-CoV-2 S retrovirus that was subsequently transduced into A549 cells (**Fig. 3a**). The S protein expression on A549 is weaker compared to 293T cells (data not shown). The pre-sorting transduction efficiency on A549 cell line was around 70% verified by flow cytometry. Transduced A549-Spike cells were subsequently sorted to achieve homogeneously high expression levels of S protein compared with 293T-hACE2-RBD cells (**Fig. 3b**) Next, we examined the activation of S309-CAR-NK-92MI cells by 293T-hACE2-RBD or A549-Spike using the conventional CD107a assay (31). Expectedly, we observed a significant increase in the surface level expression of CD107a molecules on S309-CAR-NK-92MI cells after co-culturing with susceptible 293T-hACE2-RBD or A549-Spike compared to that of parental 293T-hACE2 or A549 cells. An increase in total CD107a (percentage[%] and mean fluorescence intensity [MFI]) on S309-CAR-NK-92MI cells, compared to that of NK-92MI cells, was observed (**Fig. 3c**). In conclusion, S309-CAR-NK-92MI cells can be activated by both 293T-hACE2-RBD and A549-Spike cells.

**Figure 3:**
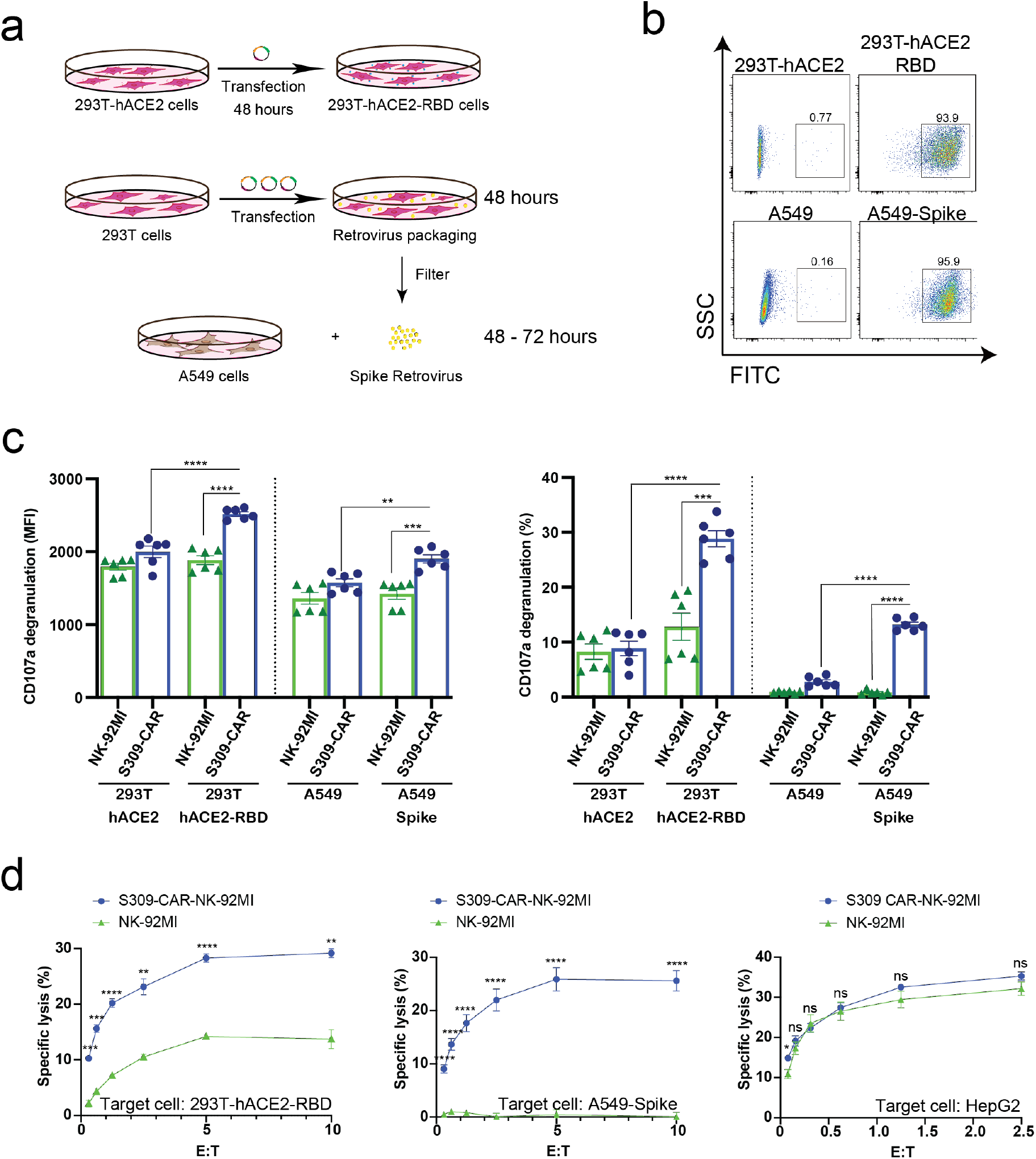
Increased CD107a surface expression and killing activity of S309-CAR-NK-92MI cells against 293T-hACE2-RBD and A549-Spike target cells. **(a)** Generation of transient 293T-hACE2-RBD and stable A549-Spike cell lines. 293T-hACE2 cells were transfected with RBD-containing plasmid for 48 hours. Transfected 293T-hACE2-RBD cells were then harvested. For the generation of A549-Spike, 293T cells were transfected with the retrovirus transfection system for 48 hours. The spike retrovirus was filtered and transduced into A549 cells for an additional 48-72 hours. (**b**) Representative dot plots showing the expressions of RBD or Spike in 293T-hACE2 or A549 cells, respectively. 293T-hACE2-RBD and A549-Spike cells were stained with anti-RBD and the expressions were confirmed by flow cytometry. The stable A549-Spike cell line was then sorted to achieve high levels of spike expression. (**c**) Quantitative data of CD107a surface expression assay of S309-CAR-NK against 293T-hACE2-RBD or A549-Spike cell lines. Briefly, S309-CAR-NK-92MI cells were cocultured with either 293T-hACE2-RBD cells, A549-Spike cells, stimulated with PMA/Ionomycin, or incubated alone for 2 hours at 37°C. Cells were then harvested and stained for CAR F(ab)2 domain [IgG (H+L)] and CD107a. Data represent mean ± SEM from two experiments. (**d**) 4-hour standard Cr^51^ release assay of S309-CAR-NK-92MI and parental NK-92MI cells against various target cell lines. 293T-hACE2-RBD, A549-Spike, and HepG2 cell lines were used as target cells for S309-CAR-NK and NK-92MI. Experimental groups were performed in triplicates. Error bars represent mean ± SEM from at least two independent experiments. Unpaired Student’s *t* test was used for both panels (**c**) and (**d**). ns p > 0.05, * p < 0.05, ** p < 0.01, *** p < 0.001, and **** p < 0.0001.

To evaluate the killing activity of S309-CAR-NK-92MI against SARS-CoV-2-protein-expressing target cells *in vitro*, we used the standard 4-hour Chromium-51 (Cr^51^) release assay (a gold standard assay to evaluate the cytotoxicity of immune cells) (32). The data show that S309-CAR-NK-92MI cells effectively kill both 293T-hACE2-RBD and A549-Spike cells by *in vitro* Cr^51^ release assay (**Fig. 3d)**. We also used an irrelevant target cell line, HepG2 (human hepatoma cell line) as a negative control, to confirm the specificity of our S309-CAR-NK. As expected, we did not observe a significant difference in the killing activity of S309-CAR-NK-92MI compared to that of wild-type NK-92MI cells (**Fig. 3d**). Here, by using three different cell lines, we demonstrated the S309-CAR-NK-92MI cells can specifically target and kill SARS-CoV-2-protein-expressing target cells.

### Expanded primary S309-CAR-NK cells can specifically kill SARS-CoV-2-protein-expressing target cells

After evaluating the killing function of S309-CAR-NK-92MI cell line, we then examined whether the expanded S309-CAR-NK^primary^ cells also have similar killing function against SARS-CoV-2-protein-expressing cells, given the natural malignancy of NK-92MI cell line(17). To expand human primary NK cells from peripheral blood (hereinafter PBNK), we isolated peripheral blood mononuclear (PBMCs) from buffy coats from healthy donors and cocultured with 100-Gy irradiated 221-mIL21 feeder cells supplemented with 200 U/mL IL-2 and 5 ng/mL IL-15, as described (33, 34). In parallel, 293T cells were transfected with a combination of plasmids containing S309-CAR in the SFG backbone, RDF, and PegPam3, as previously described (28). The SFG retrovirus particles were used to transduce expanded PBNK cells at Day 4 (**Fig. 4a**). After 48 hours, primary S309-CAR-NK cells were transferred to a G-Rex plate for continued culturing for 21 days. The NK cell purity and CAR expression were determined using flow cytometry by staining both PBNK and S309-CAR-NK^primary^ cells with anti-CD56, anti-CD3 and anti-human IgG (H+L). On average, the NK cell purity is around 90% with approximately 80% CAR transduction efficiencies for the S309-CAR-NK^primary^ (**Fig. 4b**).

**Figure 4:**
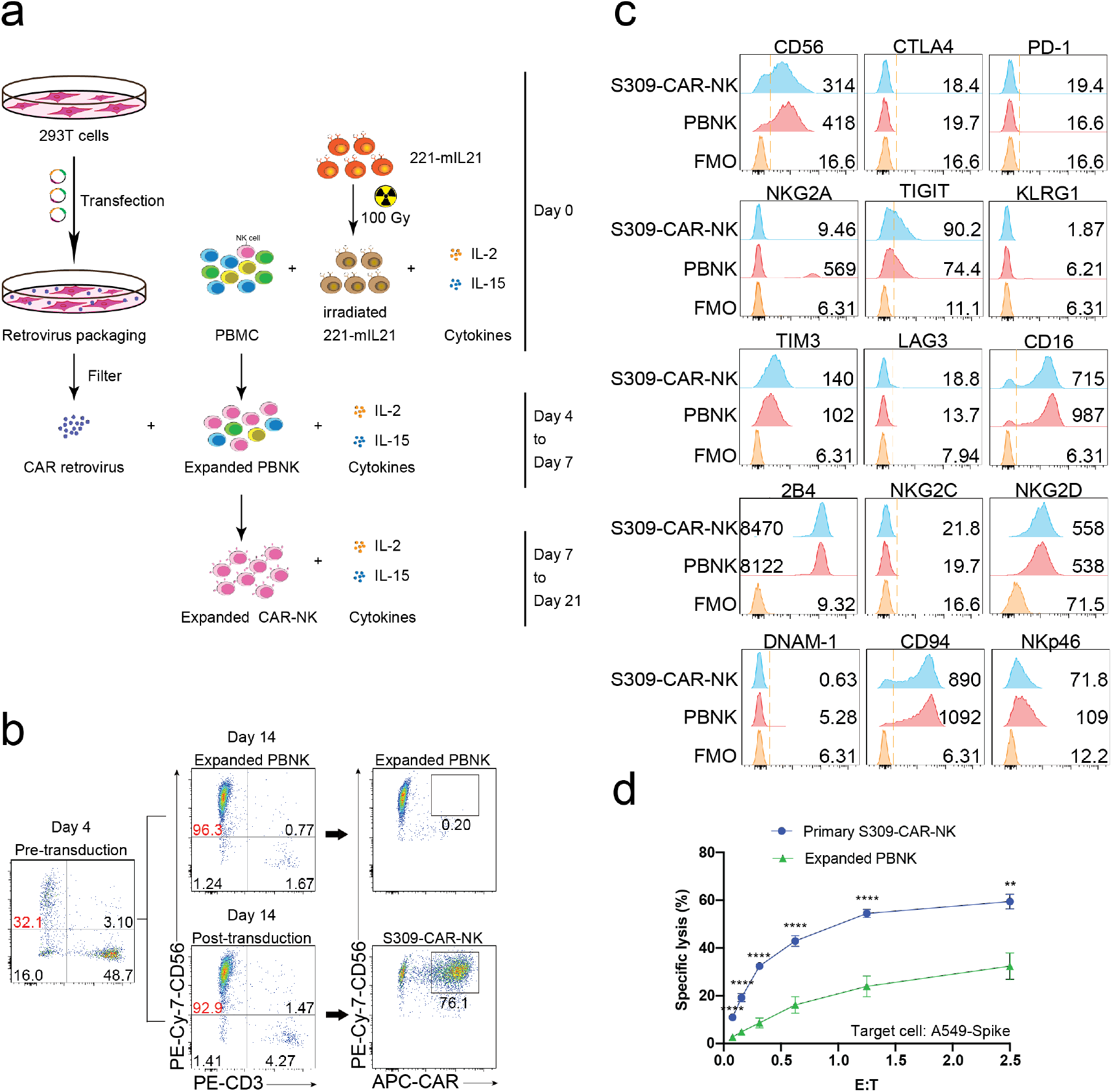
Increased killing activity of expanded S309-CAR-NK^primary^ against A549-Spike cell line. (**a**) Schematic representation of human S309-CAR-NK^primary^ expansion system. Briefly, irradiated (100 Gy) 221-mIL21 feeder cells were cocultured with PBMC supplemented with IL-2 and IL-15 on Day 0. In parallel, 293T cells were transfected with the retrovirus packaging system to produce S309-CAR retrovirus that were then transduced into the expanded PBNK cells in the presence of IL-2 and IL-15. S309-CAR-NK^primary^ cells were harvested on Day 7 and continued expansion for 21 days. (**b**) Representative dot plots of expanded primary NK cells and S309-CAR-NK^primary^. The purity of NK cells and the expression of CAR were monitored every 3-4 days. (**c**) Immunophenotyping of S309-CAR-NK^primary^ cells using flow cytometry. Antibodies against various immunomodulatory receptors including CTLA4, PD1, NKG2A, CD56, CD16, 2B4, NKG2C, NKG2D, NKp46, DNAM-1, CD94, TIGIT, KLRG1, TIM3, and LAG3 were used to stain both primary NK cells and S309-CAR-NK^primary^. (**d**) Quantitative data of cytotoxic activity of S309-CAR-NK^primary^ against A549-Spike. Briefly, expanded human S309-CAR-NK^primary^ cells were blocked with anti-CD16 for 30 minutes and then anti-NKG2D for 30 minutes on ice. The target cells were labeled with Cr^51^ for 2 hours prior to coculturing with S309-CAR-NK^primary^ cells for an additional 4 hours. The experiment was repeated at least twice. Error bars represent SEM. Unpaired Student’s *t* test was used. ** p < 0.01, *** p < 0.001, and **** p < 0.0001.

To immunophenotype the expanded primary S309-CAR-NK^primary^, we stained both expanded PBNK and S309-CAR-NK^primary^ cells for various important activating and inhibitory receptors by which the expressions were determined by flow cytometry. The inhibitory receptors include CTLA4, PD-1, NKG2A, TIGIT, KLRG1, TIM3, and LAG3. The activating receptors include CD16, 2B4, NKG2C NKG2D, DNAM-1, CD94, and NKp46 (**Fig. 4c**). In general, the expressions of these immunomodulatory receptors are comparable between expanded PBNK and S309-CAR-NK^primary^ cells.

Similar to S309-CAR-NK-92MI, we also used the 4-hour Cr^51^ release assay to evaluate the killing function of expanded primary S309-CAR-NK^primary^. The data show that S309-CAR-NK^primary^ cells effectively kill A549-Spike cells compared to expanded PBNK cells (**Fig. 4d)**. In summary, we demonstrated the expanded S309-CAR-NK^primary^ cells can also kill the SARS-CoV-2-protein-expressing target cells, indicating S309-CAR-NK^primary^ could potentially become a promising COVID-19 treatment.

### S309-CAR-NK cells have higher killing activities than CR3022-CAR-NK cells against SAR-CoV-2-protein-expressing cells

CR3022-CAR-NK-92MI cell line was previously generated (25). Previous studies showed that S309 neutralizing antibody recognizes both open and closed conformations of the SARS-CoV-2 S trimer, however, CR3022 can only bind to the open state (**Fig. 5a**) (24, 35). To further determine the functionalities of S309-CAR-NK in vitro, we compared the activation levels of S309-CAR-NK-92MI and CR3022-CAR-NK-92MI cells when cocultured with susceptible 293T-hACE2-RBD or A549-Spike target cells. As expected, we did not observe a significant difference in total CD107a, in both percentages and total mean fluorescence intensity (MFI), against the 293T-hACE2-RBD target cell. However, the expression levels of surface CD107a on CR3022-CAR-NK cells were significantly lower when cocultured with A549-Spike compared to that of S309-CAR-NK cells (**Fig. 5b**). Altogether, these data suggest that the conformation of the SARS-CoV-2 Spike trimers play a critical role in CAR recognition and binding ability and superior activation capabilities by S309-CAR molecules.

**Figure 5:**
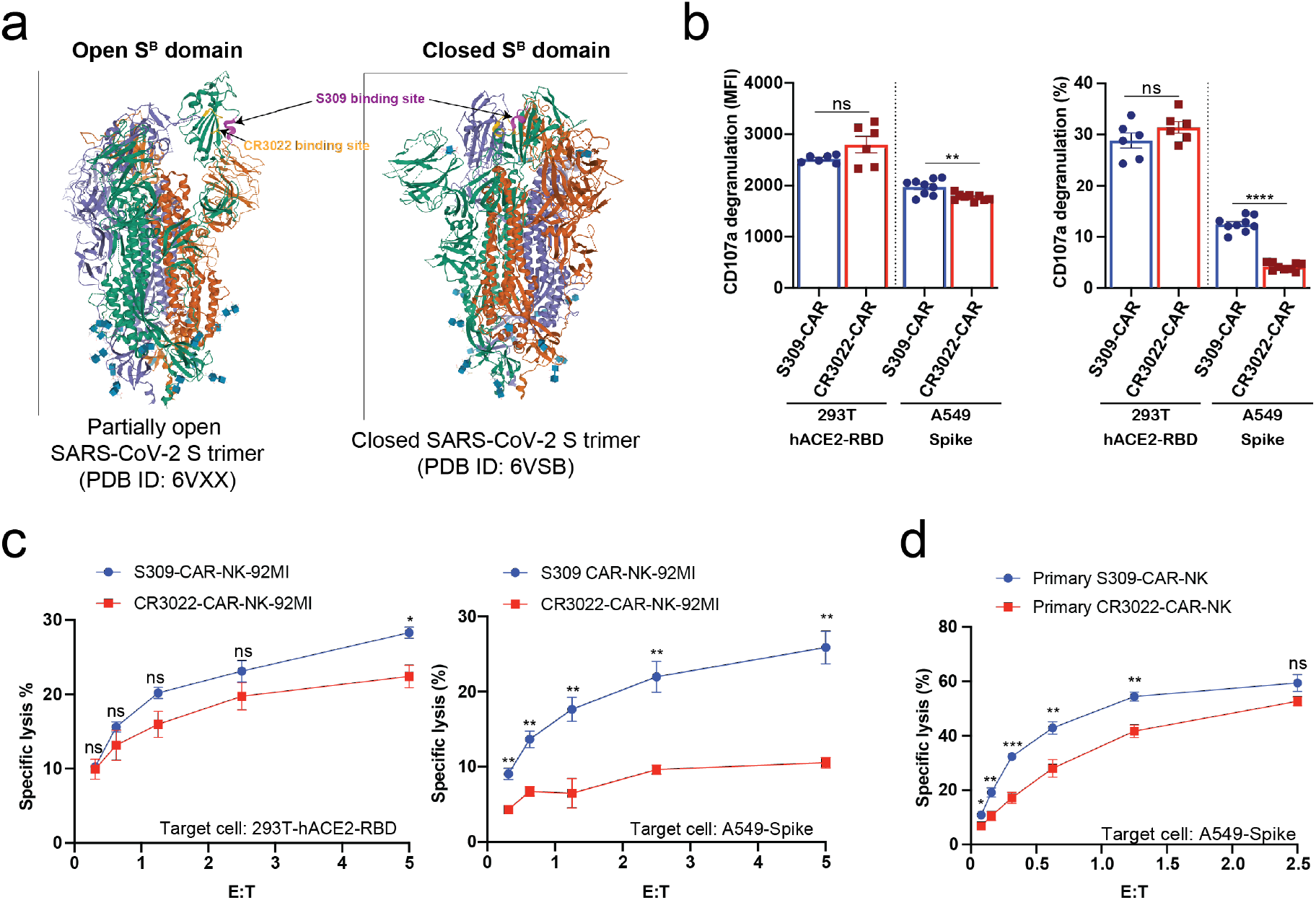
S309-CAR-NK cells are superior to CR3022-CAR-NK cells. (**a**) Diagram of S309 and CR3022 neutralizing antibodies binding to different epitopes of the SARS-CoV-2 S protein. Both open and closed conformation states of SARS-CoV-2 S protein are shown. S309 binding site is indicated in magenta and CR3022 binding site is indicated in yellow. (**b**) Quantitative data of CD107a surface expression of both S309-CAR-NK-92MI and CR3022-CAR-NK-92MI. Both transient 293T-hACE2-RBD and stable A549-Spike cell lines were used as target cells. Error bars represent SEM from at least two independent experiments. (**c**) Comparison of killing activity of S309-CAR and CR3022-CAR using the 4-hour Cr^51^ release assay. Effector cells were cocultured with Cr^51^-labeled target cells at 37°C for 4 hours. The assay was repeated for at least two times per target cell line. (**d**) Expanded S309-CAR-NK^primary^ has increased killing activity against A549-Spike cells than primary CR3022-CAR-NK^primary^. Effector cells were blocked with anti-CD16 and anti-NKG2D prior to coculturing with A549-Spike target cells for 4 hours at 37°C. Data were pooled from three independent experiments. Unpaired Student’s t test was employed for all panels. ns p > 0.05, * p < 0.05, ** p < 0.01, *** p < 0.001, and **** p<0.0001.

To further evaluate the function of S309-CAR-NK-92MI cells, we compared its killing activities to the CR3022-CAR-NK-92MI cells. Both 293T-hACE2-RBD and A549-Spike cells were used as susceptible target cells. Consistent with the results in **Fig. 5b**, we also observed a significant decrease in the killing activity of CR3022-CAR-NK cells when A549-Spike cells were the susceptible target cell line in a 4-hour Cr^51^ release assay (**Fig. 5c**). Considering NK-92MI is a cell line and may not fully reflect the true functions of primary CAR-NK cells (17), we also generated S309-CAR-NK^primary^ and CR3022-CAR-NK^primary^ from expanded PBNK. Expectedly, S309-CAR-NK^primary^ cells have better killing activities than CR3022-CAR-NK^primary^ (**Fig. 5d**), indicating that S309-CAR-NK is superior to CR3022-CAR-NK, and may have a better potential in targeting and killing SARS-CoV-2-protein-expressing cells *in vivo*.

### S309-CAR-NK^primary^ cells can also target SARS-CoV-2 D614G variant

Next, we evaluated the ability by which S309-CAR-NK^primary^ cells bind to pseudotyped SARS-CoV-2 D614G variant by flow cytometry. As expected, S309-CAR-NK^primary^ cells are unaffected by the G614 mutation and have the ability to bind to pseudotyped SARS-CoV-2 G614 variant with similar binding efficiency compared to the SARS-CoV-2 D614 **(Fig. 6a)**. To ensure the functions of S309-CAR-NK^primary^ are not altered by the G614 mutation, we also generated A549-Spike D614G cell line. Site-directed mutagenesis was performed to mutagenize SFG-SARS-CoV-2 S plasmid (**Table 1**). Subsequently, we generated A549-Spike D614G cell line using retrovirus system, which 293T cells were transduced with a combination of RDF, Pegpam3, and SARS-CoV-2 S D614G in SFG backbone. We then performed an intracellular cytokine assay to quantify the intracellular productions of TNF-α and IFN-γ of S309-CAR-NK^primary^ cells **(Fig. 6b)**. In addition to using expanded PBNK, CD19-CAR-NK cells (specific for the treatment of B-acute lymphoblastic leukemia) (21) were also included to show the specificity of S309-CAR-NK^primary^ against target cells expressing S protein or D614G variant. Consistent with our previous result in **Fig. 5d**, S309-CAR-NK^primary^ produce higher amount of intracellular TNF-α and IFN-γ compared to that of CR3022-CAR-NK^primary^ when cocultured with A549-Spike target cells. More importantly, the intracellular cytokine productions of S309-CAR-NK^primary^ cells when cocultured with A549-Spike D614G target cells were similar to that of A549-Spike, indicating S309-CAR-NK^primary^ cells can also target and kill cells infected by SARS-CoV-2 D614G variant. Surprisingly, TNF-α production slightly decreased when CR3022-CAR-NK^primary^ cells were cocultured with A549-Spike D614G but not IFN-γ **(Fig. 6b)**.

**Figure 6:**
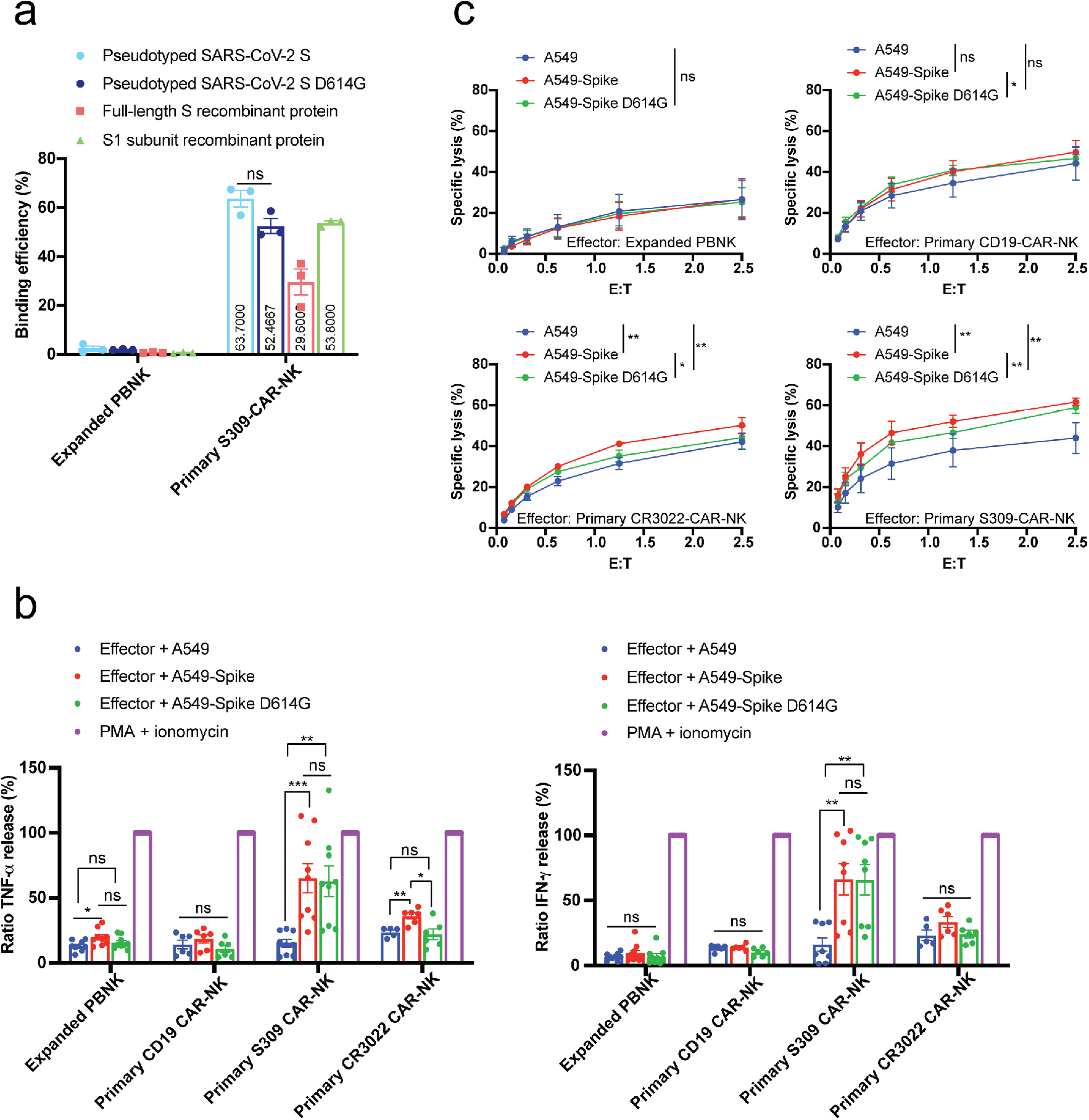
S309-CAR-NK^primary^ cells can also target SARS-CoV-2 D614G variant. **(a)** Quantitative data of the binding efficiency of S309-CAR-NK^primary^ to pseudotyped SARS-CoV-2 and D614G variant viral particles. **(b)** Intracellular cytokine productions of various effector cells. Briefly, effector cells were cocultured with A549, A549-Spike, or A549-Spike D614G for 4 hours at 37°C. Cells were then stained for surface markers and permeabilized prior to staining for intracellular cytokines. Data were acquired by flow cytometry and the values were normalized with the PMA + ionomycin group (positive control). Data were pooled from at least two independent experiments. **(c)** Expanded S309-CAR-NK^primary^ has increased killing activity against A549-Spike and A549-Spike D614G cells compared to CR3022-CAR-NK^primary^. Effector cells were blocked with anti-CD16 and anti-NKG2D prior to coculturing with A549-Spike or A549-Spike D614G target cells for 4 hours at 37°C. Data were pooled from two independent experiments. Unpaired Student’s t test was employed for panel **a**. One-way ANOVA was used for panels **b** and **c**. Post-hoc paired Student’s t test was employed after one-way ANOVA. ns p > 0.05, * p < 0.05, ** p < 0.01, *** p < 0.001, and **** p<0.0001.

We also used Cr^51^ release assay to further determine the killing activities of S309-CAR-NK^primary^ cells against A549-Spike D614G target cells. Consistently, S309-CAR-NK^primary^ cells show superior killing activities compared to CR3022-CAR-NK^primary^ in both A549-Spike and A549-Spike D614G **(Fig. 6c)**. Surprisingly, CD19-CAR-NK^primary^ cells show high cytotoxicity against A549 cell line (despite lacing expression of CD19) which may bring new findings to the field of cancer immunotherapy. Altogether, our data suggest that S309-CAR-NK^primary^ cells are able to bind the SARS-CoV-2 D614G variant in addition to killing target cells expressing S protein exhibiting the G614 mutation.

## DISCUSSION

In this study, we proposed the use of S309-CAR-NK cells for the prevention of SARS-CoV-2 infection and treatment of COVID-19 for various reasons. Firstly, S309 is a unique antibody by which it targets one of the most highly conserved epitopes in the RBD of SARS-CoV-2 and related viruses, possessing high binding affinities and neutralization potencies for both SARS-CoV-1 and SARS-CoV-2 (24). Our results suggest that this approach would retain activity against the Spike gene variants that are rapidly spreading around the globe, which may result in the development of escape mutants to the typical neutralizing antibodies in convalescent and vaccine sera. This high level of conservation suggests that S309 antibody, and the CAR-NK cells derived from this antibody, may also possess protective activity against new SARS-CoV isolates that may emerge in the future. Secondly, accumulating evidence showing NK cell involvement in the control of various viral infections (17, 21). As innate immune cells, NK cells can rapidly respond to viral infections by secreting IFN-*γ* and TNF-*α* cytokines to upregulate the defense mechanism. NK cells can also directly recognize and target viral-infected cells via antibody-dependent cellular cytotoxicity (ADCC). Thirdly, the half-life of monoclonal antibody therapeutics is approximately 21 days; however, expression of the antibody scFv portion on NK cells can prolong the protection by increasing the persistence of NK cells *in vivo*. Given these reasons, it is essential to design genetically modified NK cells that specifically target SARS-CoV-2 infected cells to control COVID-19 disease progression.

Recent clinical trials of CAR-modified NK cell-based immunotherapies have shown promising results for treating cancers and infectious diseases (17). One of the foremost challenges of using primary NK cells for immunotherapy is obtaining an adequate number of NK cells from peripheral blood or cord blood before expansion. Therefore, most of the current CAR-NK immunotherapies in clinical trials utilize NK-92 cell lines as they are easily manufactured for “off-the-shelf” purposes (34). The superior and optimized human primary NK cell expansion technology has been recently developed (33). In this study, we show the successful generation of S309-CAR-NK cells using both the NK-92MI cell line and primary NK cells expanded from human peripheral blood. S309-CAR-NK cells have the ability to efficiently bind to both SARS-CoV-2 D614 and G614 variants. We also demonstrated the cytotoxic functions of S309-CAR-NK against different target cells expressing RBD, Spike D614, and Spike G614 in various *in vitro* assays. Interestingly, we observed the superior cytotoxic functions for S309-CAR-NK compared to CR3022-CAR-NK cells against A549-Spike target cells. A plausible explanation is the different epitopes and affinities of these two neutralizing antibodies to SARS-CoV-2 S glycoprotein. CR3022 binds to a cryptic epitope that is only accessible when at least two out of the three S^B^ domains of S glycoprotein trimers are in the open conformation (35), while S309 binds to the SARS-CoV-2 S^B^ domain and to the ectodomain trimer of the S glycoprotein (24). This is reflected in neutralizing activity of S309, but not CR3022 for SARS-CoV-2, and this difference in the specific binding domains of CR3022 and S309 could explain the superior killing activities of S309-CAR-NK cells. Collectively, these preliminary data support the therapeutic potential of using S309-CAR-NK cells preventatively and as a treatment for severe COVID-19 patients. However, several limitations in the current study need to be addressed: Firstly, the pseudotyped SARS-CoV-2 S viral particles used in our experiments may not fully recapitulate the authentic SARS-CoV-2 virus. Secondly, our studies provide limited mechanistic insights showing how S309-CAR-NK cells target SARS-CoV-2 virus or SARS-CoV-2 S expressing cells. Currently, we are testing the efficacy of S309-CAR-NK cells on animal studies. Our current study supports the potential efficacy of S309-CAR-NK cells and the safety assessment of S309-CAR-NK cells *in vivo*. Future studies using wild-type SARS-CoV-2 virus in pre-clinical animal models are needed to test the efficacy and toxicity of S309-CAR-NK cells *in vivo*.

Nevertheless, the development of this novel CAR-NK cell therapy for the treatment of severe COVID-19 patients with maximal efficacy and minimal toxicity will be required to reduce patient risk and enhance the clinical benefit of these expensive and time-intensive therapies in the acute care setting. This work pioneers the use of S309-CAR-NK cells to prevent the SARS-CoV-2 infection and treat SARS-CoV-2 infected patients and will lead to the development of novel immunotherapeutic strategies for patients with immunocompromised conditions. These patients include: 1) older adults (particularly over 70-year-old with immune-compromising conditions); 2) those with comorbidities or chronic conditions such as cancer, heart disease, pre-existing lung disease, diabetes, malnutrition, and certain genetic disorders; 3) those on specific medications or treatments such as steroids, chemotherapy, radiation therapy, stem cell therapy, bone marrow or organ transplant, who may not create an effective immune protection elicited by vaccines. Furthermore, our preliminary results will expedite preclinical studies and a potential clinical application of “off-the-shelf” CAR-NK products for COVID-19 treatment. This work will also pioneer the use of a cancer immunotherapy in treating infectious diseases, broaden the use of CAR-NK cells, and initiate the development of ‘off-the-shelf’ CAR-NK products in the near future.

### Online content

Any methods, additional references, source data, and statements of code and data availability are available online.

## METHODS AND MATERIALS

### Antibodies and Reagents

PE anti-human CD3 antibody (clone OKT3), FITC and PE/Cy7 anti-human CD56 antibody (clone HCD56, BioLegend), PE anti-human CD69 antibody (clone FN50, BioLegend), PE anti-human CD8a antibody (clone RPA-T8, BioLegend), APC/Fire 750 anti-human CD226 antibody (DNAM-1) (clone 11A8, BioLegend), APC/Fire 750 anti-human KLRG1 (MAFA) antibody (clone SA231A2, BioLegend), BV421 anti-human CD335 (NKp46) antibody (clone 9E2, BioLegend), PE/Cy7 anti-human CD244 (2B4) antibody (clone C1.7, BioLegend), PE anti-human CD152 (CTLA-4) antibody (clone BNI3), APC anti-human CD366 (Tim-3) antibody (clone F38-2E2), PerCP/Cy5.5 anti-human TIGIT (VSTM3) antibody (clone A15153G), FITC anti-human CD223 (LAG-3) antibody (clone 11C3C65, BioLegend), BV510 anti-human CD314 (NKG2D) antibody (clone 1D11), APC anti-human CD94 (clone DX22, BioLegend), AF-700 anti-human IFN-γ (clone B27, BioLegend), and PE anti-human TNF-α (clone Mab11, BioLegend) were purchased from BioLegend (San Diego, CA, USA). APC anti-human CD16 antibody (clone 3G8, BD Biosciences), BV711 anti-human CD314 (NKG2D) antibody (clone 1D11, BD Biosciences), and FITC anti-human CD107a antibody (clone H4A3, BD Biosciences) were purchased from BD Biosciences (San Jose, CA, USA). PE anti-human NKG2C/CD159c antibody (clone 134591, R&D Systems) were purchased from R&D Systems. AF647 Goat anti-human IgG(H+L) F(ab’)_2_ fragment antibody was purchased from Jackson ImmunoResearch (West Grove, PA, USA). Anti-SARS-CoV-2 Coronavirus Spike protein (subunit 1) polyclonal antibody was purchased from SinoBiological (Beijing, China). Anti-SARS-CoV-2 Spike RBD rabbit polyclonal antibody was purchased from SinoBiological (Beijing, China). Anti-His mouse monoclonal antibody IgG1 (clone H-3) was purchased from Santa Cruz Biotechnology (Dallas, TX, USA). Alexa Fluor 488 goat anti-rabbit IgG (H+L) and Alexa Fluor 488 goat anti-mouse IgG1 (γ1) were purchased from Fisher Scientific (Waltham, MA).

### Cell lines

293T, A549, HepG2, and NK-92MI cell lines were purchased from the American Type Culture Collection (ATCC). 293T-hACE2 cell line is a gift from Dr. Abraham Pinter (Rutgers-New Jersey Medical School, PHRI). To maintain the stable expression of hACE2, 293T-hACE2 cells were cultured in DMEM (Corning) supplemented with 10% (v/v) fetal bovine serum (FBS), 100 U/mL Penicillin-Streptomycin (Corning), and 1μg/mL of puromycin at 37°C under 5% (v/v) CO_2_.

### Generation of transient 293T-hACE2-RBD cell line

To establish the transient 293T-hACE2-RBD cell line, 293T-hACE2 cells were transfected with 0.5 μg of SARS-CoV-2-RBD plasmid (a gift from Dr. Abraham Pinter) in each well in a 24-well plate (Eppendorf) for 48 hours at 37°C under 5% (v/v) CO_2_. Transfected cells were harvested after 48 hours and stained with primary anti-RBD (SinoBiological) followed by a goat anti-rabbit fluorophore-conjugated secondary antibody to determine the expression of RBD by flow cytometry.

### Generation of stable A549-Spike and A549-Spike D614G cell lines

pcDNA3.1-SARS-CoV-2 Spike (Addgene plasmid #145302) was used to clone SARS-CoV-2 S gene into the SFG backbone using the In-Fusion Cloning kit (Takara Bio). pSFG-SARS-CoV-2 S D614G was cloned from pSFG-SARS-CoV-2 S plasmid using the Q5 Site-Directed Mutagenesis Kit (New England BioLabs). 293T cells were transfected with 3.75 μg pSFG-SARS-CoV-2 S or pSFG-SARS-CoV-2 S D614G, 2.5 μg RDF, and 3.75 μg PegPam3 for 48 hours at 37°C under 5% (v/v) CO_2_. The spike retrovirus supernatant was filtered (0.45 μm) and transduced into A549 cells for an additional 48 – 72 hours at 37°C under 5% (v/v) CO_2_. After 2 – 3 days, transduced cells were changed to fresh DMEM (Corning) supplemented with 10% (v/v) fetal bovine serum (FBS), 100 U/mL Penicillin-Streptomycin (Corning). The spike protein expression was determined by flow cytometry by staining the transduced cells with anti-RBD antibody (SinoBiological) followed by a goat anti-rabbit fluorophore-conjugated secondary antibody. A549-Spike cells were cultured for a few days prior to sorting using anti-RBD. Sorted cells were cultured in DMEM supplemented with 10% (v/v) FBS, and 100 U/mL Penicillin-Streptomycin.

### Production of pseudotyped SARS-CoV-2 viral particles

Briefly, 293T cells were transfected using a lentivirus system with a combination of plasmids including pLP1, pLP2, pCMV-luciferase-ecoGFP (a gift from Cornell University), and pcDNA 3.1-SARS-CoV-2 Spike (Addgene plasmid #145032) for 72 hours at 37°C under 5% (v/v) CO_2_. The pseudovirus was then filtered (0.45 μm). To confirm the presence of pseudotyped SARS-CoV-2 viral particles, the filtered pseudovirus supernatant was used to transfect 293T-hACE2 for 48 hours at 37°C under 5% (v/v) CO_2_. The GFP expression of the transfected 293T-hACE2 cells was observed using an EVOS FL microscope (Life Technologies). The presence of the SARS-CoV-2 pseudovirus was further confirmed by flow cytometry, transfected 293T-hACE2 cells were stained with primary anti-RBD followed by goat anti-rabbit fluorophore-conjugated secondary antibody.

### S309-CAR construction and retrovirus production

A codon-optimized DNA fragment was synthesized by GENEWIZ encoding the S309-specific scFv and sub-cloned into the SFG retroviral vector retroviral backbone in-frame with the hinge component of human IgG1, CD28 trans-membrane domain, intracellular domain CD28 and 4-1BB, and the ζ chain of the human TCR/CD3 complex. Both the codon optimized anti-S309 scFv fragment and the SFG vector were digested with restriction endonucleases *Sal*I and *BsiW*I. SFG-S309 plasmid was transformed into Stbl3 chemically competent cells. Maxiprep was performed to enrich DNA concentration for the transfection step.

To produce S309-CAR retrovirus, 293T cells were transfected with 3.75 μg S309-CAR in SFG backbone, 3.75 μg PegPam3, and 2.5 μg RDF. S309-CAR retrovirus was harvested after 48-72 hours, filtered with a 0.45 μm filter, and transduced to NK-92MI cells in a 24-well plate coated with 0.5 µg/ml of RetroNectin diluted in PBS (Clontech). Two days later, cells were transferred to 75 cm^2^ flask (Corning) in complete NK-92MI medium (MEM-α with 12.5% (v/v) FBS, 12.5% (v/v) heat inactivated horse serum, 11 μM βME, 2 μM folic acid, and 20 μM inositol. To determine the expression of CAR or to sort S309-CAR-NK-92MI cell line, cells were stained with anti-CD56 and anti-human IgG(H+L) F(ab’)_2_ fragment.

### Primary NK cell expansion from peripheral blood

Human blood related work was approved by the Rutgers University Institutional Review Board (IRB). Lymphocyte Separation Medium (Corning) was used to isolate PBMCs from the buffy coats purchased from New York Blood Center. To expand human primary NK cells, 5 ×10^6^ cells of isolated PBMCs were cocultured with 10 ×10^6^ cells of 100 Gy-irradiated 221-mIL21 cells in 30 mL RPMI 1640 media (Corning) supplemented with 10% (v/v) FBS, 2 mM L-Glutamine (Corning), 100 U/mL Penicillin-Streptomycin, 200 U/mL IL-2 (PeproTech), and 5 ng/mL IL-15 (Peprotech) in a G-REX 6 Multi-well culture plate (Wilson Wolf) at 37°C under 5% (v/v) CO_2_. Media were changed every 3 – 4 days. An automated cell counter (Nexcelom Bioscience, Lawrence, MA, USA) was used to count the total cell numbers. The NK cell purity was determined by staining cells with anti-CD56 and anti-CD3 followed by flow cytometry analysis.

### Transduction of expanded NK cells with S309-CAR

The transduction procedure was previously described, briefly, 293T cells were transfected with a combination of SFG-S309, PegPam3, and RDF. S309-CAR retrovirus was harvested after 48-72 hours, filtered, and transduced to Day 4 of expanded primary NK cells in a 24-well plate coated with 0.5 µg/ml of RetroNectin. Transduced cells were harvested and transferred to a G-Rex well in 30 mL RPMI 1640 media supplemented with 10% (v/v) FBS, 2 mM L-Glutamine, 100 U/mL Penicillin-Streptomycin, 200 U/mL IL-2 (PeproTech), and 5 ng/mL IL-15 (Peprotech). Media were changed every 3 – 4 days up to 21 days. Cells were stained for CD56, CD3, and anti-human IgG (H+L) F(ab’)_2_ fragment for the determination of NK cell purity and CAR expression, followed by flow cytometry analysis.

### S309-CAR and RBD binding assay

To evaluate the binding activity of CR3022-CAR to RBD domain of SARS-CoV-2 S, S309-CAR or NK-92MI (5 × 10^5^) cells were incubated with 5 µg of His-gp70-RBD recombinant protein (a gift from Dr. Abraham Pinter) in DPBS buffer (0.5 mM MgCl_2_ and 0.9 mM CaCl_2_ in PBS) in for 30 minutes on ice. Cells were washed twice with PBS, stained with anti-His in FACS buffer (0.2% FBS in PBS) for 30 minutes on ice and then washed twice again with PBS. Cells were then stained with goat anti-mouse (IgG1) secondary antibody in FACS buffer for 30 minutes on ice, washed twice with PBS, and analyzed by Flow Cytometry.

### S309-CAR and pseudotyped SARS-CoV-2 S viral particles binding assay

S309-CAR, NK-92MI, and 293T-hACE2 (5 × 10^5^) cells were first equilibrated with BM binding media (complete RPMI-1640 containing 0.2% BSA and 10 mM HEPES pH 7.4). Due to the non-specific binding to the S309-CAR of our secondary antibody, cells were first blocked with anti-human IgG(H+L) F(ab’)_2_ fragment for 30 minutes on ice in BM and washed thrice with PBS. Full-length recombinant S protein (Acrobio systems), and S1 subunit recombinant protein (a gift from Dr. Abraham Pinter) were diluted with BM to appropriate concentrations. Filtered Pseudotyped SARS-CoV-2 S was used immediately following filtration without further dilution. Pseudotyped SARS-CoV-2 S, or 1 µg of full-length recombinant S protein, or 1 µg of S1 subunit recombinant protein was added to designated wells of a 96-well V bottom plate. The plate was centrifuged at 600 × g for 30 minutes at 32°C, and subsequently incubated at 37°C at 5% CO_2_ for 1 hour. Cells were washed twice with PBS, stained with anti-S1 (SinoBiological) in FACS buffer (2% FBS in PBS) for 30 minutes on ice and washed thrice with PBS. Cells were then stained with goat anti-rabbit secondary antibody in FACS buffer for 30 minutes on ice, washed thrice with PBS, and analyzed by Flow Cytometry.

### Flow cytometry analysis

Cells were stained and washed as previously described. Cells were analyzed on a FACS LSRII or an LSR Fortessa flow cytometer (36). PMT voltages were adjusted and compensation values were calculated before data collection. Data were acquired using FACS Diva software (36) and analyzed using FlowJo software (36).

### CD107a degranulation assay

The CD107a degranulation assay was described previously^3^. Briefly, NK-92MI or S309-CAR-NK-92MI or CR3022-CAR-NK-92MI cells (5 × 10^4^) were cocultured with 1 × 10^5^ 293T-hACE2, 293T-hACE2-RBD, A549, or A549-Spike cells in the presence of GolgiStop (BD Biosciences) in a V-bottomed 96-well plate in complete RPMI-1640 media at 37°C under 5% CO_2_ for 2 hours. The cells were harvested, washed, and stained for anti-CD3, anti-CD56, and anti-CD107a for 30 minutes, and analyzed by flow cytometry.

### Intracellular cytokine assay

Effector cells were cocultured with target cells in 5:1 ratio at 37°C under 5% CO_2_ for 4 hours in the presence of GolgiStop. Cells were then collected and stained for anti-CD3, anti-CD56, and anti-human IgG(H+L) F(ab’)_2_ fragment in FACS buffer for 30 minutes on ice. Cells were then permeabilized in the dark for 20 minutes on ice and were subsequently washed and stained for anti-TNF-α and anti-IFN-γ in permeabilization wash buffer for 30 minutes on ice prior to analyzing by Flow Cytometry.

### Cr^51^ release assay

To evaluate the cytotoxic activity of CAR-NK cells, the standard 4-hour Cr^51^ release assay was used. Briefly, target cells were labeled with Cr^51^ at 37°C for 2 hours and then resuspended at 1×10^5^/mL in NK-92MI culture medium with 10% FBS.

Then, 1×10^4^ target cells were incubated with serially diluted CAR-NK or NK-92MI cells at 37°C under 5% CO_2_ for 4 hours. After centrifugation, the supernatants were collected and transferred to a 96-well Luma plate and the released Cr^51^ was measured with a gamma counter (Wallac, Turku, Finland). The cytotoxicity (as a percentage) was calculated as follows: [(sample − spontaneous release) / (maximum release − spontaneous release)] × 100.

### Statistical analysis

Data were represented as mean ± SEM. The statistical significance was determined using a two-tailed unpaired Student t test, a two-tailed paired Student t test, a one-way ANOVA, where indicated. P < 0.05 was considered statistically significant.

## Figure and Figure legends

**Supplementary Table 1:**
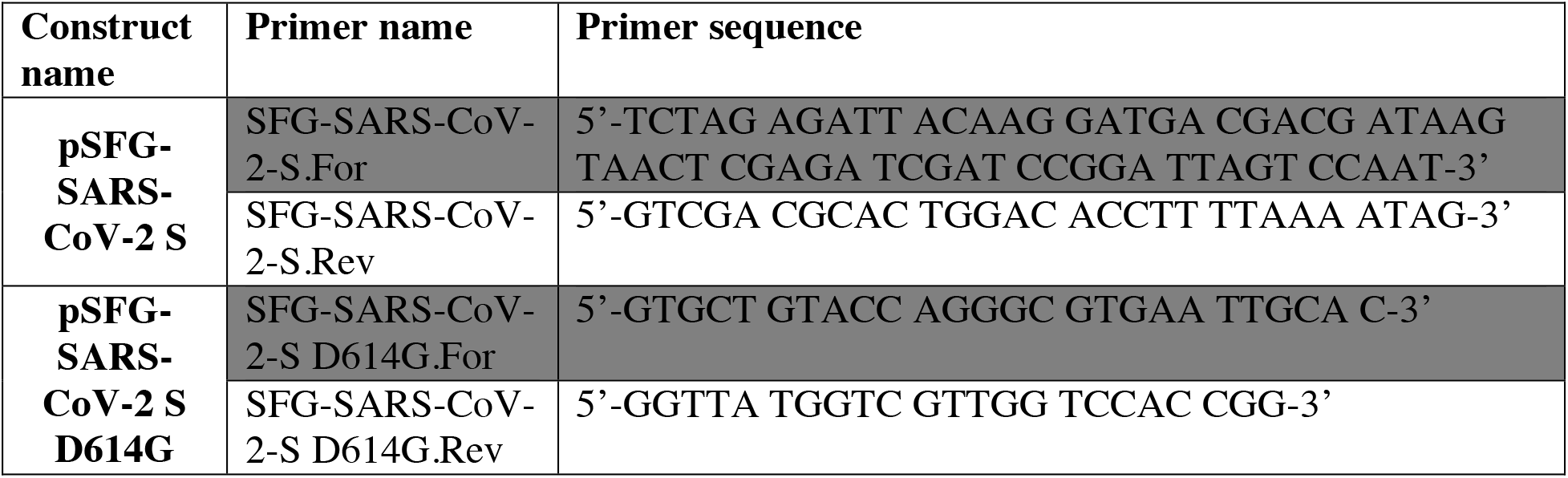
Plasmid construction.

**Supplementary Figure 1:**
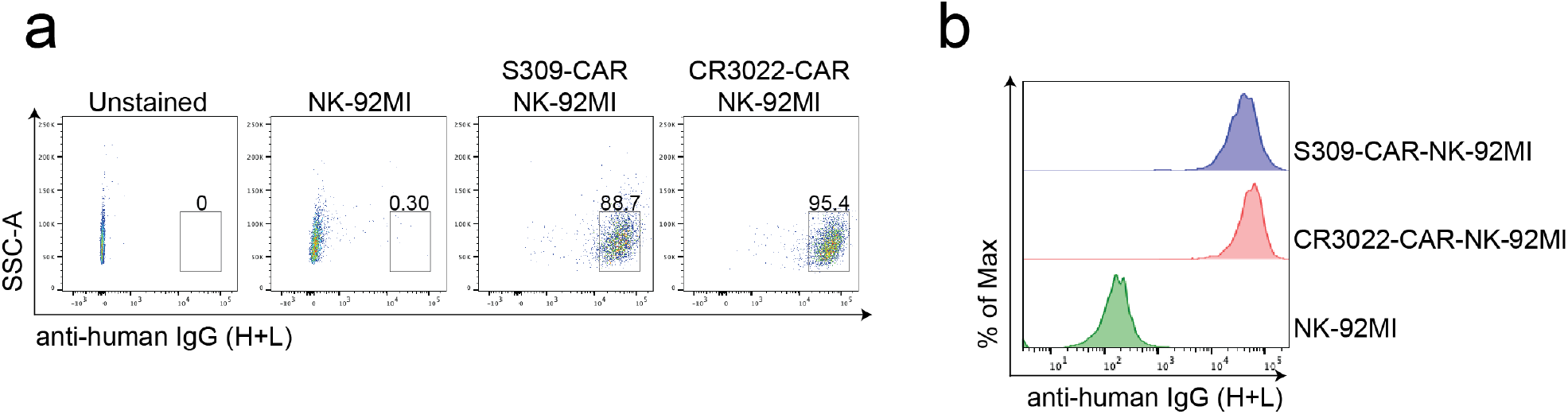
CAR expression of S309 and CR3022 in NK-92MI cell line confirmed by flow cytometry. **(a)** Flow cytometric analysis of S309-CAR-NK-92MI and CR3022-CAR-NK-92MI. Parental NK-92MI was used as a control. Cells were stained for goat anti-human IgG (H+L). The CAR expression is comparable between S309-CAR and CR3022-CAR, at 88.7% and 95.4%, respectively **(b)** Offset flow cytometric profile of S309-CAR-NK-92MI and CR3022-CAR-NK-92MI. The CAR expression is between 104 and 105 mean fluorescence intensity (MFI) for both S309-CAR and CR3022-CAR. Data are representative of at least two independent experiments.

**Supplementary Figure 2:**
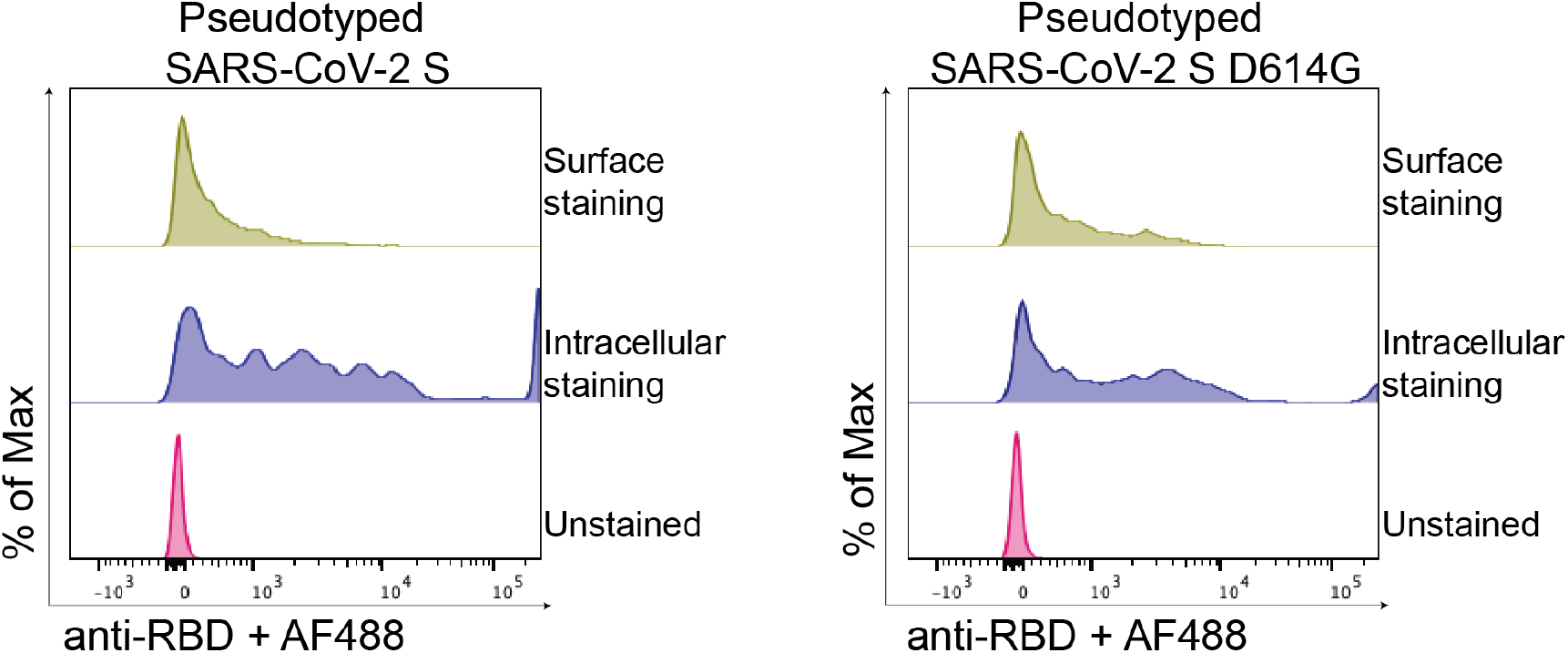
Confirmation of pseudotyped SARS-CoV-2 D614 and SARS-CoV-2 G614 viral particles production. Briefly, produced SARS-CoV-2 pseudovirus was used to infect 293T-hACE2 cells. After 48-72 hours, 293T-hACE2 cells were collected to detect SARS-CoV-2 S viral particles by flow cytometry. Intracellular or surface staining was performed with anti-RBD in permeabilization wash buffer or FACS buffer, respectively, for 30 minutes. Cells were then washed and stained for goat anti-rabbit AF488 prior to flow cytometry analysis.

## Acknowledgements

We would like to thank the members of the Liu laboratory for their comments on the manuscripts. We thank Dr. Rongfu Wang (Houston Methodist Research Institute) for providing pHAGE-FFLuc-GFP plasmid. We also thank Dr. Abraham Pinter (Rutgers-Public Health Research Institute) for providing 293T-ACE2 cell line, Dr. Pei-Hui Wang (Advanced Medical Research Institute, Cheeloo College of Medicine, Shandong University, Jinan 250012, China) and Dr. Yuan Liu (Cornell University) for providing the related SARS-CoV-2 plasmids. We also would like to thank Dr. Gianpietro Dotti (37) for the SFG vectors. This work was supported in part from HL125018 (D. Liu), AI124769 (D. Liu), AI129594 (D. Liu), AI130197 (D. Liu), and Rutgers University-New Jersey Medical School Liu Laboratory Startup funding.

## Author contributions

M.M., S.B., and D.L. designed the study and wrote the manuscript, other authors assisted with experiments and manuscript preparation. D.L. supervised the study.

## Additional information

Supplementary information is available for this paper on the Journal website. Correspondence and requests for materials should be addressed to D.L.

## References

1. R. Li, S. Pei, B. Chen, Y. Song, T. Zhang, W. Yang and J. Shaman: Substantial undocumented infection facilitates the rapid dissemination of novel coronavirus (SARS-CoV-2). Science, 368(6490), 489–493 (2020) doi:10.1126/science.abb3221

2. L. Yurkovetskiy, X. Wang, K. E. Pascal, C. Tomkins-Tinch, T. P. Nyalile, Y. Wang, A. Baum, W. E. Diehl, A. Dauphin, C. Carbone, K. Veinotte, S. B. Egri, S. F. Schaffner, J. E. Lemieux, J. B. Munro, A. Rafique, A. Barve, P. C. Sabeti, C. A. Kyratsous, N. V. Dudkina, K. Shen and J. Luban: Structural and Functional Analysis of the D614G SARS-CoV-2 Spike Protein Variant. Cell, 183(3), 739–751 e8 (2020) doi:10.1016/j.cell.2020.09.032

3. J. A. Plante, Y. Liu, J. Liu, H. Xia, B. A. Johnson, K. G. Lokugamage, X. Zhang, A. E. Muruato, J. Zou, C. R. Fontes-Garfias, D. Mirchandani, D. Scharton, J. P. Bilello, Z. Ku, Z. An, B. Kalveram, A. N. Freiberg, V. D. Menachery, X. Xie, K. S. Plante, S. C. Weaver and P. Y. Shi: Spike mutation D614G alters SARS-CoV-2 fitness. Nature (2020) doi:10.1038/s41586-020-2895-3

4. E. Volz, V. Hill, J. T. McCrone, A. Price, D. Jorgensen, A. O’Toole, J. Southgate, R. Johnson, B. Jackson, F. F. Nascimento, S. M. Rey, S. M. Nicholls, R. M. Colquhoun, A. da Silva Filipe, J. Shepherd, D. J. Pascall, R. Shah, N. Jesudason, K. Li, R. Jarrett, N. Pacchiarini, M. Bull, L. Geidelberg, I. Siveroni, C.-U. Consortium, I. Goodfellow, N. J. Loman, O. G. Pybus, D. L. Robertson, E. C. Thomson, A. Rambaut and T. R. Connor: Evaluating the Effects of SARS-CoV-2 Spike Mutation D614G on Transmissibility and Pathogenicity. Cell (2020) doi:10.1016/j.cell.2020.11.020

5. J. ter Meulen, E. N. van den Brink, L. L. M. Poon, W. E. Marissen, C. S. W. Leung, F. Cox, C. Y. Cheung, A. Q. Bakker, J. A. Bogaards, E. van Deventer, W. Preiser, H. W. Doerr, V. T. Chow, J. de Kruif, J. S. M. Peiris and J. Goudsmit: Human monoclonal antibody combination against SARS coronavirus: Synergy and coverage of escape mutants. Plos Medicine, 3(7), 1071–1079 (2006) doi:ARTN E237 10.1371/journal.pmed.0030237

6. A. Cizmecioglu, H. A. Cizmecioglu, M. H. Goktepe, A. Emsen, C. Korkmaz, F. E. Tasbent, F. Colkesen and H. Artac: Apoptosis-Induced T Cell Lymphopenia Is Related to COVID-19 Severity. J Med Virol (2020) doi:10.1002/jmv.26742

7. S. Varchetta, D. Mele, B. Oliviero, S. Mantovani, S. Ludovisi, A. Cerino, R. Bruno, A. Castelli, M. Mosconi, M. Vecchia, S. Roda, M. Sachs, C. Klersy and M. U. Mondelli: Unique immunological profile in patients with COVID-19. Cell Mol Immunol (2020) doi:10.1038/s41423-020-00557-9

8. J. W. Song, C. Zhang, X. Fan, F. P. Meng, Z. Xu, P. Xia, W. J. Cao, T. Yang, X. P. Dai, S. Y. Wang, R. N. Xu, T. J. Jiang, W. G. Li, D. W. Zhang, P. Zhao, M. Shi, C. Agrati, G. Ippolito, M. Maeurer, A. Zumla, F. S. Wang and J. Y. Zhang: Immunological and inflammatory profiles in mild and severe cases of COVID-19. Nat Commun, 11(1), 3410 (2020) doi:10.1038/s41467-020-17240-2

9. S. Jamwal, A. Gautam, J. Elsworth, M. Kumar, R. Chawla and P. Kumar: An updated insight into the molecular pathogenesis, secondary complications and potential therapeutics of COVID-19 pandemic. Life Sciences, 257 (2020) doi:ARTN 118105 10.1016/j.lfs.2020.118105

10. F. N. Release: FDA Takes Additional Action in Fight Against COVID-19 By Issuing Emergency Use Authorization for Second COVID-19 Vaccine. In, fda.gov (2020)

11. F. N. Release: FDA Takes Key Action in Fight Against COVID-19 By Issuing Emergency Use Authorization for First COVID-19 Vaccine. In, (2020)

12. E. E. Walsh, R. W. Frenck, Jr., A. R. Falsey, N. Kitchin, J. Absalon, A. Gurtman, S. Lockhart, K. Neuzil, M. J. Mulligan, R. Bailey, K. A. Swanson, P. Li, K. Koury, W. Kalina, D. Cooper, C. Fontes-Garfias, P. Y. Shi, O. Tureci, K. R. Tompkins, K. E. Lyke, V. Raabe, P. R. Dormitzer, K. U. Jansen, U. Sahin and W. C. Gruber: Safety and Immunogenicity of Two RNA-Based Covid-19 Vaccine Candidates. N Engl J Med, 383(25), 2439–2450 (2020) doi:10.1056/NEJMoa2027906

13. F. P. Polack, S. J. Thomas, N. Kitchin, J. Absalon, A. Gurtman, S. Lockhart, J. L. Perez, G. Perez Marc, E. D. Moreira, C. Zerbini, R. Bailey, K. A. Swanson, S. Roychoudhury, K. Koury, P. Li, W. V. Kalina, D. Cooper, R. W. Frenck, Jr., L. L. Hammitt, O. Tureci, H. Nell, A. Schaefer, S. Unal, D. B. Tresnan, S. Mather, P. R. Dormitzer, U. Sahin, K. U. Jansen, W. C. Gruber and C. C. T. Group: Safety and Efficacy of the BNT162b2 mRNA Covid-19 Vaccine. N Engl J Med (2020) doi:10.1056/NEJMoa2034577

14. A. T. Widge, N. G. Rouphael, L. A. Jackson, E. J. Anderson, P. C. Roberts, M. Makhene, J. D. Chappell, M. R. Denison, L. J. Stevens, A. J. Pruijssers, A. B. McDermott, B. Flach, B. C. Lin, N. A. Doria-Rose, S. O’Dell, S. D. Schmidt, K. M. Neuzil, H. Bennett, B. Leav, M. Makowski, J. Albert, K. Cross, V. V. Edara, K. Floyd, M. S. Suthar, W. Buchanan, C. J. Luke, J. E. Ledgerwood, J. R. Mascola, B. S. Graham, J. H. Beigel and R. N. A. S. G. m: Durability of Responses after SARS-CoV-2 mRNA-1273 Vaccination. N Engl J Med (2020) doi:10.1056/NEJMc2032195

15. A. Curti, L. Ruggeri, A. D’Addio, A. Bontadini, E. Dan, M. R. Motta, S. Trabanelli, V. Giudice, E. Urbani, G. Martinelli, S. Paolini, F. Fruet, A. Isidori, S. Parisi, G. Bandini, M. Baccarani, A. Velardi and R. M. Lemoli: Successful transfer of alloreactive haploidentical KIR ligand-mismatched natural killer cells after infusion in elderly high risk acute myeloid leukemia patients. Blood, 118(12), 3273–3279 (2011) doi:10.1182/blood-2011-01-329508

16. J. S. Miller, Y. Soignier, A. Panoskaltsis-Mortari, S. A. McNearney, G. H. Yun, S. K. Fautsch, D. McKenna, C. Le, T. E. Defor, L. J. Burns, P. J. Orchard, B. R. Blazar, J. E. Wagner, A. Slungaard, D. J. Weisdorf, I. J. Okazaki and P. B. McGlave: Successful adoptive transfer and in vivo expansion of human haploidentical NK cells in patients with cancer. Blood, 105(8), 3051–3057 (2005) doi:10.1182/blood-2004-07-2974

17. D. Liu, S. Tian, K. Zhang, W. Xiong, N. M. Lubaki, Z. Chen and W. Han: Chimeric antigen receptor (CAR)-modified natural killer cell-based immunotherapy and immunological synapse formation in cancer and HIV. Protein Cell, 8(12), 861–877 (2017) doi:10.1007/s13238-017-0415-5

18. D. Liu, S. Badeti, G. Dotti, J. G. Jiang, H. Wang, J. Dermody, P. Soteropoulos, D. Streck, R. B. Birge and C. Liu: The Role of Immunological Synapse in Predicting the Efficacy of Chimeric Antigen Receptor (CAR) Immunotherapy. Cell Commun Signal, 18(1), 134 (2020) doi:10.1186/s12964-020-00617-7

19. N. Shah, L. Li, J. McCarty, I. Kaur, E. Yvon, H. Shaim, M. Muftuoglu, E. Liu, R. Z. Orlowski, L. Cooper, D. Lee, S. Parmar, K. Cao, C. Sobieiski, R. Saliba, C. Hosing, S. Ahmed, Y. Nieto, Q. Bashir, K. Patel, C. Bollard, M. Qazilbash, R. Champlin, K. Rezvani and E. J. Shpall: Phase I study of cord blood-derived natural killer cells combined with autologous stem cell transplantation in multiple myeloma. Br J Haematol, 177(3), 457–466 (2017) doi:10.1111/bjh.14570

20. D. F. Liu, S. Tian, K. Zhang, W. Xiong, N. Lubaki, Z. Y. Chen and W. D. Han: Chimeric antigen receptor (CAR)-modified natural killer cell-based immunotherapy and immunological synapse formation in cancer and HIV. Protein & Cell, 8(12), 861–877 (2017) doi:10.1007/s13238-017-0415-5

21. E. L. Liu, D. Marin, P. Banerjee, H. A. Macapinlac, P. Thompson, R. Basar, L. N. Kerbauy, B. Overman, P. Thall, M. Kaplan, V. Nandivada, I. Kaur, A. N. Cortes, K. Cao, M. Daher, C. Hosing, E. N. Cohen, P. Kebriaei, R. Mehta, S. Neelapu, Y. Nieto, M. Wang, W. Wierda, M. Keating, R. Champlin, E. J. Shpall and K. Rezvani: Use of CAR-Transduced Natural Killer Cells in CD19-Positive Lymphoid Tumors. New England Journal of Medicine, 382(6), 545–553 (2020) doi:10.1056/NEJMoa1910607

22. P. Zhou, X. L. Yang, X. G. Wang, B. Hu, L. Zhang, W. Zhang, H. R. Si, Y. Zhu, B. Li, C. L. Huang, H. D. Chen, J. Chen, Y. Luo, H. Guo, R. D. Jiang, M. Q. Liu, Y. Chen, X. R. Shen, X. Wang, X. S. Zheng, K. Zhao, Q. J. Chen, F. Deng, L. L. Liu, B. Yan, F. X. Zhan, Y. Y. Wang, G. F. Xiao and Z. L. Shi: A pneumonia outbreak associated with a new coronavirus of probable bat origin. Nature, 579(7798), 270–273 (2020) doi:10.1038/s41586-020-2012-7

23. W. H. Li, M. J. Moore, N. Vasilieva, J. H. Sui, S. K. Wong, M. A. Berne, M. Somasundaran, J. L. Sullivan, K. Luzuriaga, T. C. Greenough, H. Choe and M. Farzan: Angiotensin-converting enzyme 2 is a functional receptor for the SARS coronavirus. Nature, 426(6965), 450–454 (2003) doi:10.1038/nature02145

24. D. Pinto, Y. J. Park, M. Beltramello, A. C. Walls, M. A. Tortorici, S. Bianchi, S. Jaconi, K. Culap, F. Zatta, A. De Marco, A. Peter, B. Guarino, R. Spreafico, E. Cameroni, J. B. Case, R. E. Chen, C. Havenar-Daughton, G. Snell, A. Telenti, H. W. Virgin, A. Lanzavecchia, M. S. Diamond, K. Fink, D. Veesler and D. Corti: Cross-neutralization of SARS-CoV-2 by a human monoclonal SARS-CoV antibody. Nature, 583(7815), 290–295 (2020) doi:10.1038/s41586-020-2349-y

25. M. Ma, S. Badeti, K. Geng and D. Liu: Efficacy of Targeting SARS-CoV-2 by CAR-NK Cells. bioRxiv (2020) doi:10.1101/2020.08.11.247320

26. D. J. Giard, S. A. Aaronson, G. J. Todaro, P. Arnstein, J. H. Kersey, H. Dosik and W. P. Parks: In vitro cultivation of human tumors: establishment of cell lines derived from a series of solid tumors. J Natl Cancer Inst, 51(5), 1417–23 (1973) doi:10.1093/jnci/51.5.1417

27. R. T. Eastman, J. S. Roth, K. R. Brimacombe, A. Simeonov, M. Shen, S. Patnaik and M. D. Hall: Remdesivir: A Review of Its Discovery and Development Leading to Emergency Use Authorization for Treatment of COVID-19. Acs Central Science, 6(5), 672–683 (2020) doi:10.1021/acscentsci.0c00489

28. W. Xiong, Y. Chen, X. Kang, Z. Chen, P. Zheng, Y. H. Hsu, J. H. Jang, L. Qin, H. Liu, G. Dotti and D. Liu: Immunological Synapse Predicts Effectiveness of Chimeric Antigen Receptor Cells. Mol Ther, 26(4), 963–975 (2018) doi:10.1016/j.ymthe.2018.01.020

29. X. Tian, C. Li, A. Huang, S. Xia, S. Lu, Z. Shi, L. Lu, S. Jiang, Z. Yang, Y. Wu and T. Ying: Potent binding of 2019 novel coronavirus spike protein by a SARS coronavirus-specific human monoclonal antibody. Emerg Microbes Infect, 9(1), 382–385 (2020) doi:10.1080/22221751.2020.1729069

30. J. Lan, J. Ge, J. Yu, S. Shan, H. Zhou, S. Fan, Q. Zhang, X. Shi, Q. Wang, L. Zhang and X. Wang: Structure of the SARS-CoV-2 spike receptor-binding domain bound to the ACE2 receptor. Nature, 581(7807), 215–220 (2020) doi:10.1038/s41586-020-2180-5

31. D. Liu, Y. T. Bryceson, T. Meckel, G. Vasiliver-Shamis, M. L. Dustin and E. O. Long: Integrin-dependent organization and bidirectional vesicular traffic at cytotoxic immune synapses. Immunity, 31(1), 99–109 (2009) doi:10.1016/j.immuni.2009.05.009

32. J. L. McCoy, R. B. Herberman, E. B. Rosenberg, F. C. Donnelly, P. H. Levine and C. Alford: 51 Chromium-release assay for cell-mediated cytotoxicity of human leukemia and lymphoid tissue-culture cells. Natl Cancer Inst Monogr, 37, 59–67 (1973)

33. Y. Yang, S. Badeti, H. C. Tseng, M. T. Ma, T. Liu, J. G. Jiang, C. Liu and D. Liu: Superior Expansion and Cytotoxicity of Human Primary NK and CAR-NK Cells from Various Sources via Enriched Metabolic Pathways. Mol Ther Methods Clin Dev, 18, 428–445 (2020) doi:10.1016/j.omtm.2020.06.014

34. J. H. Gong, G. Maki and H. G. Klingemann: Characterization of a human cell line (NK-92) with phenotypical and functional characteristics of activated natural killer cells. Leukemia, 8(4), 652–8 (1994)

35. M. Yuan, N. C. Wu, X. Zhu, C. D. Lee, R. T. Y. So, H. Lv, C. K. P. Mok and I. A. Wilson: A highly conserved cryptic epitope in the receptor binding domains of SARS-CoV-2 and SARS-CoV. Science, 368(6491), 630–633 (2020) doi:10.1126/science.abb7269

36. R. Romee, M. Rosario, M. M. Berrien-Elliott, J. A. Wagner, B. A. Jewell, T. Schappe, J. W. Leong, S. Abdel-Latif, S. E. Schneider, S. Willey, C. C. Neal, L. Yu, S. T. Oh, Y. S. Lee, A. Mulder, F. Claas, M. A. Cooper and T. A. Fehniger: Cytokine-induced memory-like natural killer cells exhibit enhanced responses against myeloid leukemia. Sci Transl Med, 8(357), 357ra123 (2016) doi:10.1126/scitranslmed.aaf2341

37. A. Churojana, I. Sakarunchai, T. Aurboonyawat, E. Chankaew, P. Withayasuk and B. Sangpetngam: Efficiency of endovascular therapy for bilateral cavernous sinus dural arteriovenous fistula. World Neurosurg (2020) doi:10.1016/j.wneu.2020.10.001

